# Transcriptome-wide transmission disequilibrium analysis identifies novel risk genes for autism spectrum disorder

**DOI:** 10.1101/835678

**Authors:** Kunling Huang, Yuchang Wu, Junha Shin, Ye Zheng, Alireza Fotuhi Siahpirani, Yupei Lin, Zheng Ni, Jiawen Chen, Jing You, Sunduz Keles, Daifeng Wang, Sushmita Roy, Qiongshi Lu

## Abstract

Recent advances in consortium-scale genome-wide association studies (GWAS) have highlighted the involvement of common genetic variants in autism spectrum disorder (ASD), but our understanding of their etiologic roles, especially the interplay with rare variants, is incomplete. In this work, we introduce an analytical framework to quantify the transmission disequilibrium of genetically regulated gene expression from parents to offspring. We applied this framework to conduct a transcriptome-wide association study (TWAS) on 7,805 ASD proband-parent trios, and replicated our findings using 35,740 independent samples. We identified 31 associations at the transcriptome-wide significance level. In particular, we identified *POU3F2* (p=2.1e-7), a transcription factor (TF) mainly expressed in developmental brain. TF targets regulated by *POU3F2* showed a 2.1-fold enrichment for known ASD genes (p=4.6e-5) and a 2.7-fold enrichment for loss-of-function *de novo* mutations in ASD probands (p=7.1e-5). These results provide a clear example of the connection between ASD genes affected by very rare mutations and an unlinked key regulator affected by common genetic variations.

## Introduction

Autism spectrum disorder (ASD) is a highly heritable neurodevelopmental disorder affecting 1.5% of the world population^1^. It manifests as impaired social interaction and communication, repetitive behavior, and restricted interests with highly heterogenous clinical presentations^2^. Whole-exome sequencing (WES) studies for ASD have identified numerous ultra-rare or *de novo* single-nucleotide variants, small insertions and deletions (indels), and copy number variants (CNVs)^3–7^. Although these protein-disrupting genetic variations have large effects on the disease risk, they are only found in a moderate proportion of ASD probands. It has been estimated that the contribution of *de novo* loss-of-function mutations and CNVs to the variance in ASD liability was only 3% while common genetic variants explain 50% of the variance in the population^8^. Recently, genome-wide association studies (GWAS) with large sample sizes, coupled with novel statistical genetic approaches, have provided new insights into the involvement of common single-nucleotide polymorphisms (SNPs) in ASD. Polygenic risk of ASD is significantly over-transmitted from parents to ASD probands but not their unaffected siblings in simplex families^9^. Such over-transmission was also observed in probands with *de novo* mutations in known ASD genes. Additionally, a recent GWAS meta-analysis of 18,381 ASD cases and 27,969 controls identified multiple genome-wide significant loci, but did not implicate apparent associations at ASD risk genes identified in WES studies^10^. These results suggested that distinct mechanistic pathways may underlie the ASD risk attributed to rare and common genetic variants, but our understanding of their interplay remains incomplete.

One potential approach to better dissect the genetic basis of ASD is to fine-map candidate genes affected by common SNPs and then investigate how they interact with genes harboring rare pathogenic variants implicated in WES studies. Transcriptome-wide association study (TWAS) is an analytical strategy that integrates expression quantitative trait loci (eQTL) annotations with GWAS data to identify disease genes^11–13^. Through advanced predictive modeling for gene expression traits, TWAS effectively combines association evidence across many eQTL in diverse tissues and has identified risk genes for numerous complex diseases^14^.

In this study, we introduce TITANS (TrIo-based Transcriptome-wide AssociatioN Study), a novel statistical framework to conduct TWAS in proband-parent trios. Combining recent advances in TWAS modeling and the trio-based study design in ASD cohorts, we demonstrate transmission disequilibrium of genetically regulated gene expression in brain tissues from parents to ASD probands. Specifically, we conducted GWAS and TWAS on 7,805 ASD trios from the Autism Genome Project (AGP), the Simons Simplex Collection (SSC), and the Simons Foundation Powering Autism Research for Knowledge (SPARK) cohort, and replicated our findings in an independent cohort of 13,076 cases and 22,664 controls (**Methods**). We identified 31 associations at the transcriptome-wide significance level. In particular, we identified *POU3F2*, a master regulator highly expressed in developmental brain whose downstream target genes are strongly enriched for known ASD genes and mutations.

## Results

### Transmission disequilibrium of polygenic risk, gene expression, and SNP alleles

We applied multiple analytical approaches to dissect common SNPs’ contributions to ASD risk at different scales. First, we performed polygenic transmission disequilibrium test (pTDT)^9^ to examine the transmission disequilibrium of ASD polygenic risk in probands. ASD polygenic risk scores (PRS) were constructed using case-control samples from the iPSYCH cohort (N=35,740; **Methods**). We confirmed a highly significant over-transmission of ASD PRS from parents to probands in multiple datasets (p=1.4e-25 in the meta-analysis), including the SPARK cohort which has not been previously analyzed (p=1.0e-11; **Supplementary Figure 1**). No significant over-transmission was identified in 3,245 healthy siblings (p=0.88).

Next, using a novel approach called TITANS (**Methods**), we performed a TWAS with eQTL and splicing quantitative trait loci (sQTL) in 12 brain tissues from the Genotype-Tissue Expression (GTEx) project^15^ and the CommonMind consortium (CMC)^16^. For each proband, we generated 3 pseudo siblings using phased genotype data of the parents (**Figure 1A**). We imputed gene expression and intron usage values^17^ for all probands and pseudo siblings (**Figure 1B**) using UTMOST^12^ (10 GTEx brain tissues) and FUSION^11^ (CMC dorsolateral prefrontal cortex; DLPFC) imputation models. We used conditional logistic regression^18^ to assess the transmission disequilibrium of imputed gene expression traits while adjusting for the genetic similarity between probandw and pseudo siblings. We also used the same framework to perform trio-based GWAS (**Figure 1C; Methods**).

**Figure 1.**
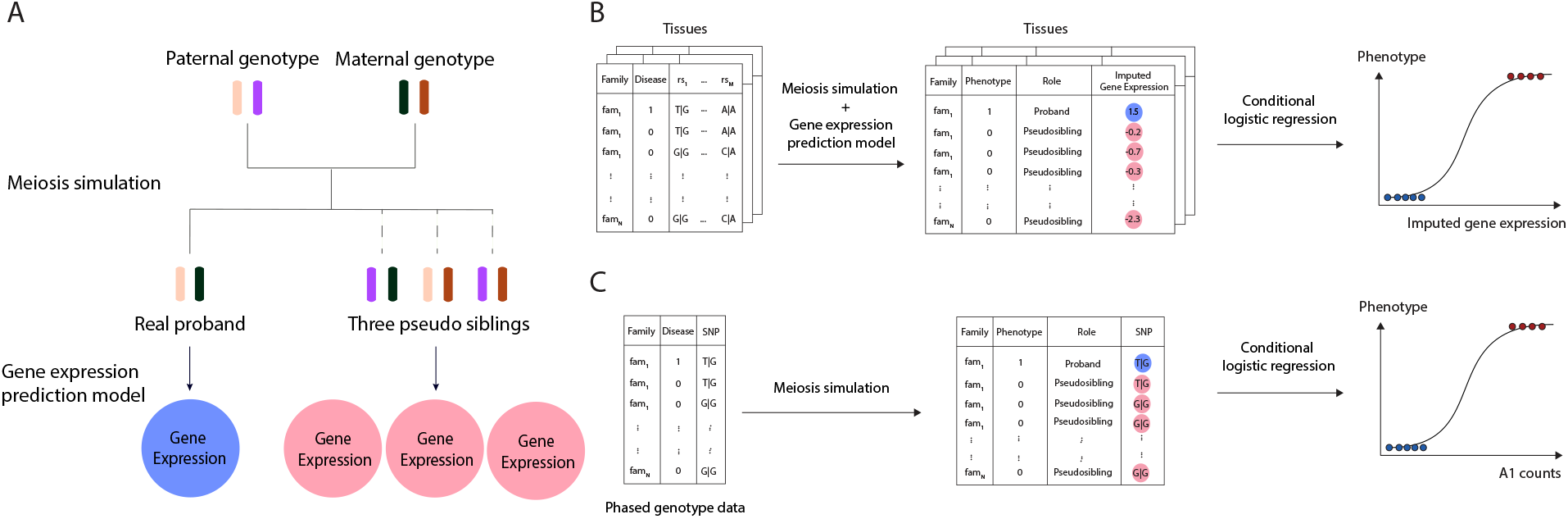
TITANS workflow. **(A)** We generate three matched pseudo siblings for each proband using the phased genotype data of parents and impute gene expression values. **(B)** We compare the impute gene expression traits between probands and matched pseudosiblings and use conditional logistic regression to quantify the associations. **(C)** We simulate genotype data for matched pseudosiblings and use conditional logistic regression to assess SNP-disease associations.

We identified significant transmission disequilibrium of *POU3F2* expression (p=5.6e-7; GTEx hippocampus) and *MSRA* intron usage (p=2.3e-7; CMC DLPFC splicing) in 7,805 trios after correcting for the number of genes in each tissue (**Table 1**). Both associations were replicated in an independent cohort of 13,076 cases and 22,664 controls (p=0.015 and 0.002, respectively). Meta-analysis enhanced the associations at *POU3F2* and *MSRA* and identified 29 additional significant associations at the transcriptome-wide significance level (**Supplementary Table 1**; **Supplementary Figures 2-11)**. 5 associations, i.e. *POU3F2* (p=2.1e-7), *MSRA* (p=5.7e-9), *MAPT* (p=3.6e-7), *KIZ* (p=1.9e-7), and *NKX2-2* (p=1.5e-10), remained significant after a stringent Bonferroni correction for all genes and all tissues in the analysis (**Table 1** and **Figure 2**). In total, these associations implicated 18 unique candidate genes from 7 loci, including 5 novel loci not previously identified in GWAS. No significant associations were identified in unaffected sibling-parent trios (**Supplementary Figure 12**) or after randomly shuffling probands and pseudo siblings (**Supplementary Figure 13**).

**Table 1.** Cross-tissue significant associations in TWAS. Beta and SE indicate the normalized effect size estimates and standard error in conditional logistic regression. Some effect size estimates are unavailable in the replication cohort since FUSION does not provide effect size estimates.

| Gene | Chr | Tissue | Discovery Stage (N=7,805 trios) |  |  | Replication Stage (N=35,740) |  |  | Meta-analysis |  |  |
| --- | --- | --- | --- | --- | --- | --- | --- | --- | --- | --- | --- |
|  |  |  | Beta | SE | P | Beta | SE | P | Beta | SE | P |
| *POU3F2* | 6 | GTEx hippocampus | 0.09 | 0.02 | 5.56E-07 | 0.03 | 0.01 | 0.015 | 0.05 | 0.01 | 2.05E-07 |
| *MSRA* | 8 | CMC DLPFC - splicing | 0.09 | 0.02 | 2.26E-07 | - | - | 0.002 | - | - | 5.67E-09 |
| *MAPT* | 17 | CMC DLPFC - splicing | 0.06 | 0.02 | 2.42E-04 | - | - | 4.09E-04 | - | - | 3.62E-07 |
| *KIZ* | 20 | CMC DLPFC | 0.05 | 0.02 | 1.73E-03 | - | - | 2.62E-05 | - | - | 1.88E-07 |
| *NKX2-2* | 20 | GTEx nucleus accumbens basal ganglia | -0.05 | 0.02 | 2.44E-03 | -0.07 | -0.01 | 2.91E-09 | -0.06 | 0.01 | 1.49E-10 |

**Figure 2.**
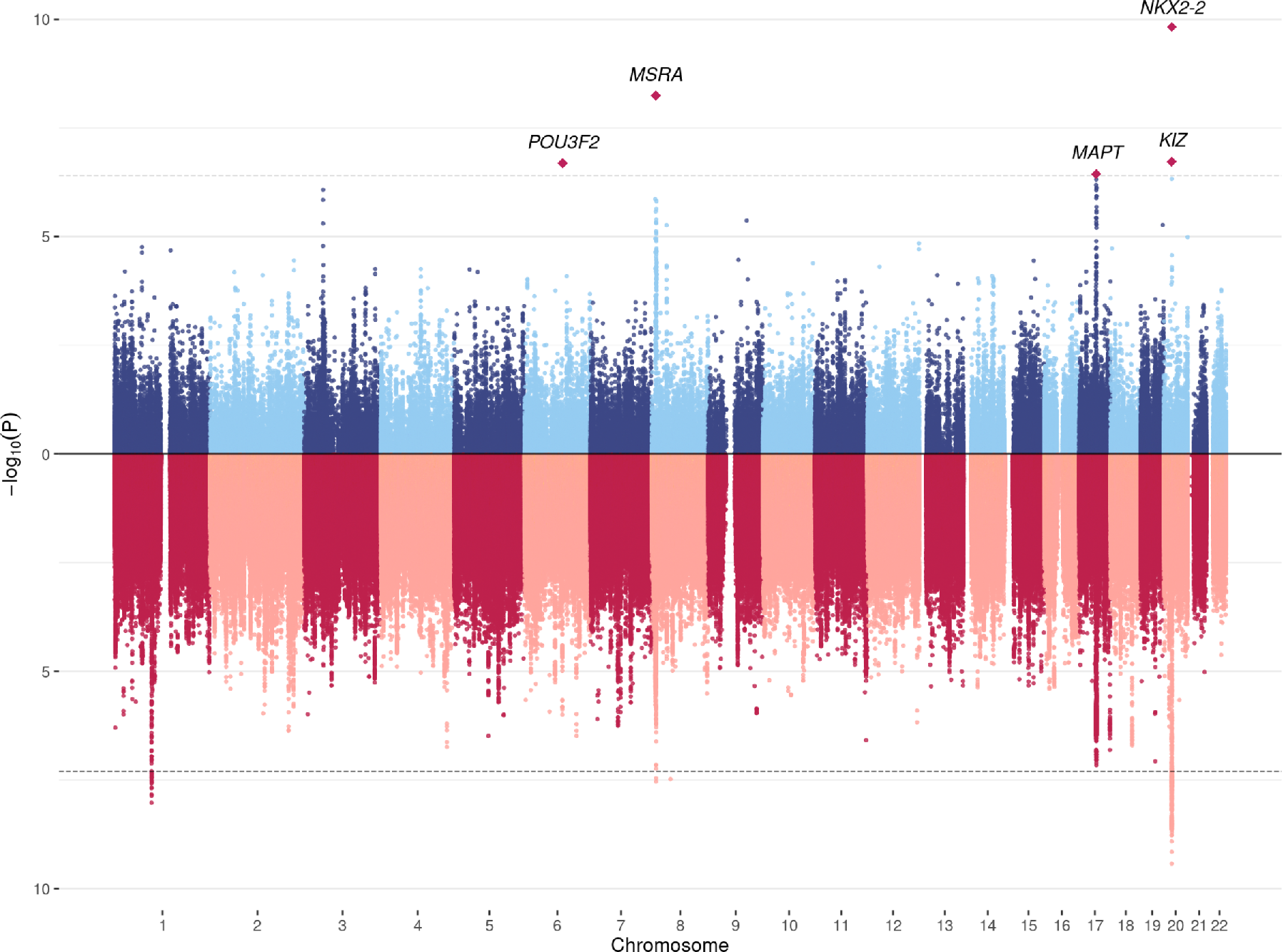
Mirrored Manhattan plot for TWAS and GWAS results. TWAS results are shown in the upper panel. GWAS associations are shown in the lower panel. The dashed line in the upper panel indicates the cross-tissue transcriptome-wide significance cutoff (p=4.0e-7) and the dashed line in the lower panel is the genome-wide significance cutoff (p=5.0e-8). TWAS associations for all 12 tissues are shown.

GWAS meta-analysis of trios and case-control cohorts identified 4 genome-wide significant loci (**Supplementary Table 2**), 3 of which (1p21.3, 8p23.1, and 20p11.23) were among previously identified loci^10^. A locus on chromosome 8 is novel but we note that the top SNP did not exist in the trio-based analysis. Overall, association patterns in GWAS and TWAS were concordant (**Figure 2**). Two GWAS loci on chromosomes 8 and 20 were also identified in TWAS. No significant associations were found in sibling-parent trios (**Supplementary Figure 12**).

### Candidate risk genes and gene set enrichment analysis

Among the 5 significant genes after a stringent Bonferroni correction for all genes and all tissues in the analysis (**Figure 3** and **Supplementary Figure 14**), *POU3F2* (also known as *BRN2*) is primarily expressed in the central nervous system (**Supplementary Figure 15**). It encodes a transcription factor with important roles in neurogenesis and brain development^19,20^. It is a known risk gene for bipolar disorder^21,22^ and has been identified as a master regulator of gene expression changes in schizophrenia and bipolar disorder^20,23^. Deletions resulting in loss of one copy of *POU3F2* cause a disorder of variable developmental delay, intellectual disability, and susceptibility to obesity^24^. Heterozygous *POU3F2* knockout mice showed deficits in adult social behavior^25^ and it has been linked to neural proliferation phenotypes in stem cell models of ASD^26^. Although this locus did not reach genome-significance in the GWAS, gene-level association at *POU3F2* was supported by a SNP-level association peak 700 kb upstream of *POU3F2* (**Figure 3A**; lead SNP rs2388334, p=1.0e-6).

**Figure 3.**
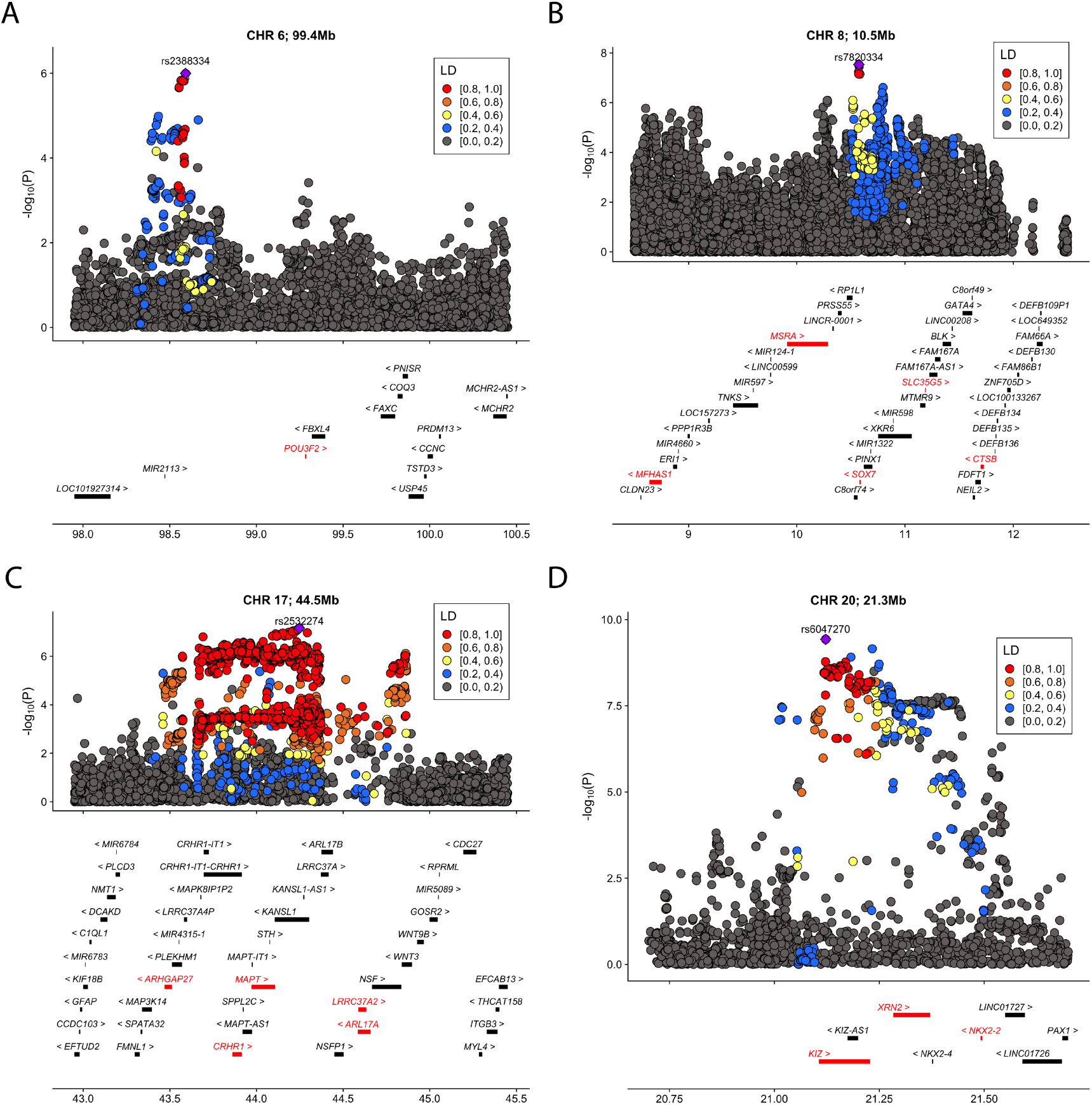
Significant loci identified in TWAS. We identified 5 cross-tissue transcriptome-wide significant associations from 4 loci. **(A)** chr1, 99.4 mb **(B)** chr8, 10.5 Mb **(C)** chr17, 44.5 mb **(D)** chr20, 21.3 mb. For each locus, the index SNP with the most significant association in GWAS is marked as purple diamond and the color of data points indicates linkage disequilibrium (LD) of neighboring SNPs with the index SNP. Genes are highlighted in red if they reached transcriptome-wide significance in at least one tissue. The x-axis denotes genome coordinates and the y-axis denotes association p-values in GWAS.

*MAPT* encodes the microtubule-associated protein tau known to associate with multiple neurodegenerative diseases including Alzheimer’s disease and Parkinson’s disease^27^ and balance of *MAPT* isoforms is critical for neuronal normal functioning^28^. This locus showed suggestive associations in the GWAS (lead SNP rs2532274, p=6.9e-8). *KIZ*, *NKX2-2*, and *MSRA* are located at 2 loci previously identified in ASD GWAS^10^. *KIZ* encodes the Kizuna centrosomal protein which is critical for stabilizing mature centrosomes during spindle formation^29^. *NKX2-2* encodes the homeobox protein NKX2.2, a transcription factor with an essential role in interpreting graded Sonic hedgehog signals and selecting neuronal identity^30^. *MSRA* shows high levels of expression in the human central nervous system and *Msra* knockout mice show abnormal behaviors^31,32^.

We investigated if genes with nominal associations (p<0.05) in TWAS are enriched in known ASD pathways. Among the 15 gene sets we tested (**Methods**), only genes encoding postsynaptic density proteins (PSD; enrichment=1.18, p=3.6e-5) and SFARI genes with evidence score 4-6 (enrichment=1.27, p=8.8e-5) showed significant enrichment for TWAS findings after multiple testing correction (**Figure 4A**; **Supplementary Table 3**). Additionally, we note that some genes with weaker evidence in the SFARI Gene database (see URLs) were identified using samples from the AGP and SSC cohorts and thus may not represent independent evidence. Notably, gene sets that are known to harbor significant burden of rare or *de novo* variants in ASD, e.g. *FMRP* target genes (enrichment=1.07, P=0.14), SFARI genes with evidence score S-3 (enrichment=1.07, p=0.24), and chromatin modifier genes (enrichment=0.94, p=0.77), showed negligible enrichment for TWAS associations. These results confirmed the distinct etiologic pathways underlying common and rare genetic variations in ASD.

**Figure 4.**
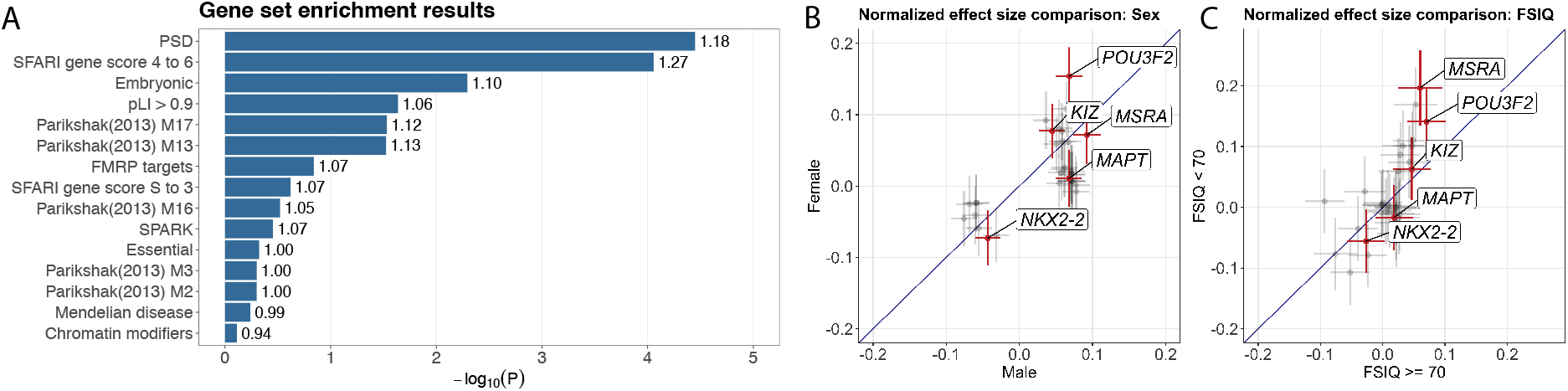
Gene set enrichment analysis and subgroup TWAS results. **(A)** Enrichment −log p-values for different gene sets are shown in the bar plot. Fold enrichment values are labeled next to each bar. **(B)** The normalized effect size estimates in sex-stratified TWAS. Effects of 31 associations identified in the pooled TWAS are shown in the plot. 5 cross-tissue significant associations are highlighted in red. For each cross, the interval indicates normalized effect ± standard error. **(C)** The normalized effect size estimates in FSIQ-stratified TWAS. Each interval indicates normalized effect ± standard error.

### TWAS associations in subgroups

Further, we investigated if the effects of candidate genes are consistent in different phenotypic subgroups. We applied TITANS to assess the 31 associations identified in TWAS in sample subgroups stratified by sex and full-scale intelligence quotient (FSIQ)^7,9^. In sex-stratified analysis of 6,484 male probands and 1,321 female probands, most genes showed comparable effect sizes in males and females (correlation=0.65; **Figure 4B**). Cross-tissue significant genes *POU3F2*, *KIZ*, and *NKX2-2* had higher effects in females. Of note, *POU3F2* showed a 2.26-fold ratio between its effects in females and in males, reaching statistical significance even under a substantially smaller sample size of female probands (**Supplementary Table 4**). This is consistent with a female protection mechanism that requires a larger effect size and risk load. We next performed FSIQ-stratified analysis and compared the transmission disequilibrium in probands with higher (FSIQ >= 70, N=2,127) and lower FSIQ (FSIQ < 70, N=731). The effect size estimates in two subgroups were mostly consistent (correlation=0.71; **Figure 4C**). *POU3F2* showed a stronger effect in the subgroup with lower FSIQ (fold=2.00; p=0.023 in subgroup with higher FSIQ, p=0.009 in subgroup with lower FSIQ).

### Regulatory role of *POU3F2* in ASD

The transcription factor encoded by *POU3F2* is a key regulator in multiple psychiatric disorders^20,23^. Based on its robust association with ASD in our analysis, we hypothesize that *POU3F2* may also play a central role in ASD through its regulatory network. We investigated the biological underpinnings of *POU3F2* by leveraging diverse types of genomic data. First, we confirmed the link between the gene-level association at *POU3F2* and GWAS associations in the same region through integrating fetal brain Hi-C data from the germinal zone (GZ) and postmitotic-zone cortical plate (CP)^33^. *POU3F2* and the GWAS association peak 700 kb upstream are located in the same topological associating domain (TAD) that is conserved in both GZ and CP zones (chr6: 97.52-99.76 mb; **Figure 5A**). Additionally, we identified 59 non-overlapping bins, each of 10 kb in size and within 1 mb from the transcription start site of *POU3F2*, showing significant interactions with the promoter region of *POU3F2* (p<1.0e-4; **Methods**; **Supplementary Tables 5-7**). Multiple bins showing significant interactions with *POU3F2* promoter colocalized with GWAS associations in this region. For example, SNP rs62422661 (p=2.0e-5 in GWAS) is located in the bin located at 98.54-98.55 mb on chromosome 6 which significantly interacts with *POU3F2* in the CP zone (p=2.0e-12). In addition, 15 SNP predictors for *POU3F2* expression, including 2 strong predictors with effect sizes ranked at top 15%, are located in bins interacting with *POU3F2* promoter (**Figure 5A**).

**Figure 5.**
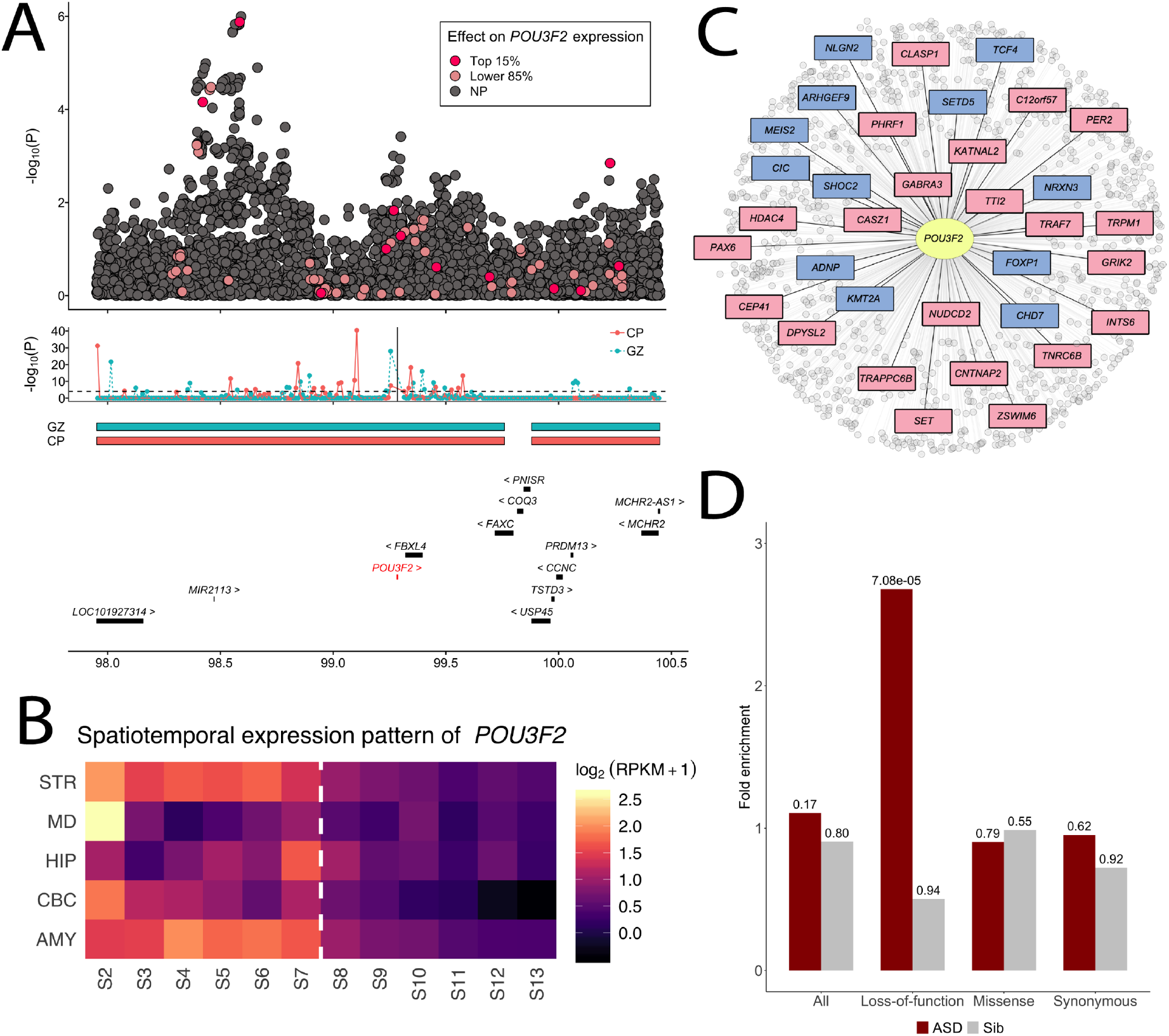
Biological underpinnings of *POU3F2*. **(A)** The upper panel shows GWAS associations at the *POU3F2* locus. Predictor SNPs in the *POU3F2* imputation model highlighted in red or pink based on their effect size rankings (top 15% or lower 85%). The middle panel shows the TADs in CP and GZ zones and the Hi-C interactions between each 10-kb bin in the region and *POU3F2* promoter which is indicated by the vertical line. The lower panel lists the genes at this locus. **(B)** The spatiotemporal expression pattern of *POU3F2* in 12 developmental stages across 5 brain regions. The periods span fetal development, infancy, childhood, adolescence, and adulthood. Average log_2_(RPKM+1) values for samples of the same region and developmental stage are shown. The dashed line indicates the boundary between later fetal and early infancy stages (0 month). **(C)** Transcription factor target genes of *POU3F2*. ASD genes in the SPARK gene list are highlighted in blue and additional genes with SFARI evidence score S to 3 are highlighted in pink. **(D)** Enrichment of *De novo* mutations in *POU3F2* targets. Enrichment results in 2,508 ASD probands and 1,911 unaffected siblings across four annotation categories (all mutations, loss-of-function, missense, and synonymous) are shown. P-values are shown above each bar.

Next, we examined the spatiotemporal expression pattern of *POU3F2* in 5 brain regions, i.e. cerebellar cortex (CBC), striatum (STR), hippocampus (HIP), mediodorsal nucleus of thalamus (MD), and amygdala (AMY), spanning from fetal development to adulthood^34^ (**Methods**). *POU3F2* showed significantly elevated expression in developmental brains compared to postnatal brains across all 5 brain regions (p=5.3e-3, permutation test; **Figure 5B**). A similar pattern was also observed in several other genes (e.g. *MAPT*) while *NKX2-2* showed elevated expression in postnatal brains (**Supplementary Figure 16**).

Additionally, we used the regulatory network from Chasman et al.^35^ to investigate the enrichment of known ASD genes in target genes regulated by *POU3F2*. The transcription factor target network of *POU3F2* contained 1,013 genes (**Figure 5C**). These genes showed strong enrichment (enrichment=2.1, p=0.012) for the SPARK genes which included 153 curated genes known to be associated with autism (**Methods**; URLs) and for SFARI genes with scores S to 3 (enrichment=2.1, p=4.6e-05). Various gene sets previously shown to enrich for rare and *de novo* mutations in ASD, including chromatin modifiers (p=2.6e-4), *FMRP* targets (p=0.009), and loss-of-function intolerant genes (p=2.2e-6), were significant enriched in *POU3F2* targets (**Supplementary Table 8**). Furthermore, *POU3F2* target genes were significantly enriched for loss-of-function *de novo* mutations (enrichment=2.68, p=7.1e-05, Poisson test; **Methods**) in 2,508 SSC probands (**Figure 5D**, **Supplementary Table 9**). Enrichment remained substantial even after we removed known ASD genes in the SPARK gene list from the analysis (enrichment=1.66, p=0.06) despite the reduced statistical evidence (**Supplementary Table 10**). We did not observe significant enrichment for missense or synonymous mutations. No enrichment was observed in 1,911 unaffected siblings.

Finally, we obtained transcription factor binding sites (TFBS) of *POU3F2* based on the prior network in Chasman et al.^35^ and used linkage disequilibrium score regression (LDSC) to assess the enrichment of ASD heritability in these TFBS^36^ (**Methods**). SNPs located near POU3F2 binding sites explained 11.7% of ASD heritability, showing a 5.3-fold enrichment with suggestive statistical significance (p=0.054; **Supplementary Table 11**).

## Discussion

In this study, we have presented TITANS, an analytical framework for testing the transmission disequilibrium of genetically regulated molecular traits between parents and probands. Through integrative modeling of GWAS data in trios and rich QTL annotations from large consortia such as GTEx^15^, this approach effectively combines association evidence at multiple SNPs to implicate risk genes affected by common genetic variations. Applied to multiple large-scale ASD cohorts including the SPARK study which has not been previously reported, we conducted a TWAS on 7,805 proband-parent trios and replicated our findings in 35,740 case-control samples. Meta-analysis identified a total of 31 transcriptome-wide significant associations, many of which are located at novel loci not previously implicated in GWAS.

Among the identified associations, convergent evidence suggested a critical etiologic role of *POU3F2* in ASD. *POU3F2* encodes a transcription factor mainly expressed in the central nervous system^19^ and has known key regulatory roles in schizophrenia and bipolar disorder^20,23^. In our analysis, it reached transcriptome-wide statistical significance in trio-based TWAS and was successfully replicated in the case-control replication. Meta-analysis strengthened the association at *POU3F2* and it remained significant after a stringent multiple testing correction for all genes and all tissues analyzed in this study. Subtype analysis suggested that *POU3F2* has enhanced over-transmission in female probands (2.3-fold) and individuals with lower FSIQ (2-fold). Furthermore, we demonstrated its etiologic importance and its connection to other ASD risk genes through integrative analysis of diverse types of genomic data. Analysis of fetal brain Hi-C data confirmed significant interactions between *POU3F2* promoter and multiple genome regions near GWAS associations located in the same TAD. Analysis of spatiotemporal gene expression data suggested significantly elevated *POU3F2* expression in developmental brain. TFBS of *POU3F2* were enriched for ASD heritability. Downstream target genes regulated by *POU3F2* were enriched for known ASD risk genes identified in WES studies. *POU3F2* targets were also significantly enriched for loss-of-function *de novo* mutations in ASD probands. Enrichment remained substantial even after known ASD genes were removed from the gene set.

WES studies have identified numerous extremely rare, protein-disrupting variants in ASD and have implicated risk genes and pathways^3–7^. Successful studies focusing on other types of genetic variants using GWAS and whole-genome sequencing approaches have just begun to emerge^9,10,37–39^. A common and somewhat puzzling observation in these studies was that common SNPs associated with ASD did not influence the same genes and pathways enriched for rare variants. Our analysis partly confirmed this observation – genes showing strong associations in TWAS had limited overlap with genes identified through WES. However, the *POU3F2* results provide a clear example of the direct link of genes affected by very rare mutations with common genetic variations at a second, unlinked locus. These findings provide insights into the interplay of common and rare genetic variations in ASD, shed light on regulatory network-based modeling of epistatic interactions, and have broad implications for the genetic basis of other diseases.

## Methods

### Sample information and data processing

We accessed AGP samples through dbGaP (accession: phs000267). The total sample size was 7,880. Genotyping was performed using the Illumina Human 1M-single Infinium BeadChip. Details on these samples have been described elsewhere (see URLs)^40^. We accessed samples from the SSC and the SPARK study through the Simons Foundation Autism Research Initiative (SFARI; see URLs). The SSC cohort contains comprehensive genotype and phenotype information from 2,600 simplex families, each family has one ASD child, and healthy parents and siblings. Genotyping was performed in batches by the Illumina IMv1, IMv3 Duo, and Omni2.5 arrays. Details on these data can be found on the SFARI website (URLs) and have been described elsewhere^39^. Samples in the SPARK study were genotyped by the Illumina Infinium Global Screening Array. Details on these samples have been previously reported^41,42^ and are available on the SFARI website (URLs)

We performed pre-imputation quality control (QC) using PLINK^43^. Only individuals with self-reported European ancestries were included in the study. SNPs with genotype call rate < 0.95, minor allele frequency (MAF) less than 0.01, or significant deviation from Hardy-Weinberg equilibrium (p<1.0e-6) were removed from the analysis. Samples with genotype missing rate > 0.05 were also excluded from the analysis. We used genetic relationship coefficients estimated from GCTA^44^ to identify and remove overlapped samples among different cohorts. After QC, 2,188, 1,794, and 3,823 independent proband-parent trios remained in AGP, SSC, and SPARK cohorts respectively. 1,432 and 1,813 trios of sibling-parent trios remained in SSC and SPARK. The UCSC liftOver tool was used to liftover the genome coordinates in AGP samples from hg18 to hg19. The genotype data were phased and imputed to the HRC reference panel version r1.1 2016 using the Michigan Imputation server^45^. We removed SNPs with imputation quality < 0.8 or MAF < 0.01 in the post-imputation QC. 7,260,224 SNPs remained in the AGP study after QC. 7,298,961 SNPs, 7,029,817 SNPs, and 6,866,248 SNPs remained in the SSC 1Mv1, 1Mv3, and Omni2.5 datasets, respectively. 7,031,717 SNPs remained in the SPARK data.

We used case-control samples from the iPSYCH cohort as the replication dataset in our study (13,076 cases and 22,664 controls). The iPSYCH ASD sample contains all Danish children born between 1981 and 2005 and details on this cohort are described elsewhere^46^. This cohort has been included in a recent ASD GWAS meta-analysis^10^. Samples in the iPSYCH cohort are independent from samples in the AGP, SSC, and SPARK.

### Polygenic transmission disequilibrium analysis

We used the iPSYCH GWAS summary statistics to generate ASD PRS on all samples. We performed a LD-clumping using PLINK with a p-value threshold of 1, a LD threshold of 0.1, and a distance threshold of 1,000 Kb. After clumping, 167085 SNPs remained in the dataset. PRSice was used for PRS calculation^47^. We quantified the transmission disequilibrium of ASD PRS using the pTDT approach^9^.

### Trio-based TWAS and GWAS analysis

We developed a statistical framework TITANS to perform trio-based TWAS (**Figure 1**). We used UTMOST^12^ gene expression imputation models for 10 brain tissues in GTEx and imputation models for CMC DLPFC expression and intron usage values implemented in FUSION^11^. UTMOST model uses a cross-tissue penalized regression model to borrow information from tissues with larger sample size and improve imputation accuracy of gene expression^12^. FUSION trains multiple imputation models in each tissue separately, including Bayesian sparse linear mixed model, elastic net, LASSO, and an ordinary least square model using single best eQTL as the predictor. We selected the best model using the cross-validation.

Given a gene with *m* predictor SNPs, we extracted those SNPs from parents’ phased genotypes and recombined the chromosomes based on Mendelian inheritance to create the genotypes of pseudo siblings. Since only cis-regulators within the local region are included in gene expression and intron usage imputation models, we assumed no crossover events in our analysis. Given the parental data, four recombined pseudo offspring genotypes can be created, each having a paternal haplotype and a maternal haplotype. We imputed gene expression and intron usage on each proband and all four simulated pseudo siblings. We excluded the pseudo sibling whose imputed expression is the closest to the proband’s since one of the four simulated offsprings’ genotype should be identical to the proband if there is no phasing error or crossover. We tested the association between imputed gene expression and disease phenotype using conditional logistic regression^18^, with conditional likelihood

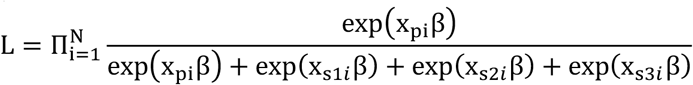

Here, *x*_*pi*_, *x*_*s1i*_, …, *x*_*s3i*_ denote the imputed gene expression or intron usage values of the proband and 3 pseudo siblings in the i^th^ family. We used the clogit function in the R package ‘survival’ to numerically estimate the effect size β, which can be interpreted as transmission disequilibrium of imputed expression. The SE of β, the z-score test statistic, and association p-value are also reported. TWAS was conducted in the AGP, SSC, and SPARK cohorts separately. Results in different trio-based cohorts were meta-analyzed using the inverse-variance weighted method^48^. These results were then meta-analyzed with the associations in the replication stage using z-score-based meta-analysis weighted by sample sizes^48^.

We performed TWAS in sample subgroups based on sex and FSIQ (**Supplementary Table 4**). We conducted sex-stratified TWAS in each cohort and meta-analyzed the result across AGP, SSC, and SPARK using the inverse-variance weighted method^48^. FSIQ-stratified analysis based on a cutoff of 70 was conducted in SSC and SPARK separately and then combined through meta-analysis.

We used a similar framework to conduct GWAS in trios. For each SNP, we create four recombined genotypes based on parental data, exclude a genotype identical to the proband’s genotype, and perform conditional logistic regression to assess the association between each SNP and ASD status.

### Gene set enrichment analysis

We used hypergeometric test to assess if genes with nominal TWAS associations (p<0.05 in any tissue) were enriched in gene sets that have bene linked to ASD in past literatures (**Supplementary Table 3**). Gene sets assessed in our analysis included co-expression modules M2, M3, M13, M16, and M17 from Parikshak et al.^49^. *FMRP* targets, genes encoding postsynaptic density proteins (PSD), gene preferentially expressed in human embryonic brains downloaded from BRAINSPAN (see URLs), essential genes^50^, chromatin modifier genes^5^, and genes with probability of loss-of-function intolerance (pLI) > 0.9 from the Exome Aggregation Consortium^51^. In addition, we downloaded genes from the SFARI Gene database on August 2019 (URLs) and created two gene sets based on evidence scores. The gene set based on scores S, 1, 2, or 3 include genes involved in ASD with high to suggestive evidence and genes predisposing to ASD in the context of a syndromic disorder. Genes with scores 4-6 have limited evidence or have only been hypothesized to link to ASD. Finally, we obtained a list of 153 genes with known roles in ASD curated by the SPARK study (URLs). We refer to this gene set of SPARK genes in our analyses.

### Hi-C analysis

We used the human fetal brain Hi-C data (URLs; GEO: GSE77565)^33^ at resolution 10 kb in the analysis. The samples were sequenced using Illumina HiSeq 2000 chip, collecting from three individuals aging gestation week (GW) 17–18 (one sample from GW17 and two samples from GW18). The Hi-C libraries were constructed in two brain zones GZ and CP. The TAD region of GZ and CP are also provided. We converted the Hi-C contact matrices (HDF5 format) normalized by ICE^52^ into the sparse contact matrix format (BED format) and leveraged Fit-Hi-C^53^ to detect the significant interactions in the regions of interest. Benjamini-Hochberg procedure^54^ was employed to control the false discovery rate.

### Spatiotemporal expression analysis

We obtained spatiotemporal gene expression data from BRAINSPAN for 17 candidate genes (URLs) with significant associations in our TWAS analysis. Average log_2_(RPKM+1) values for samples of the same region and developmental stage were calculated. Expression data were from 5 brain regions, i.e. CBC, STR, HIP, MD, and AMY, and spanned from 8 weeks post-conception (PCW) to 40 years as indicated in Kang et al^55^. mRNA sequencing was performed using the Illumina Genome Analyzer IIx. Details on these data are described elsewhere^34^.

### *POU3F2* transcription factor binding network

The transcriptional targets of *POU3F2* were obtained using the procedure from Chasman et al^35^. We downloaded *POU3F2* motif position weight matrices (PWM) from 3 databases, CIS-BP^56^, ENCODE^57^, and JASPAR^58^. We obtained DNase-I seq data for neural progenitor cells from the Roadmap Epigenome Consortium^59^ (GEO: GSE18927). Next, we applied the Protein Interaction Quantification (PIQ) algorithm^60^ to identify *POU3F2* motif binding sites across the human genome. Using the DNase-I seq data, the PIQ algorithm defines a purity score (0.5-1.0) for a motif instance, which quantifies the likelihood of a true binding event in that site. PIQ motif instances were mapped to the transcription start sites from Gencode v10 within a 10kb radius. The confidence of the edge between a transcription factor and the target was defined as the maximum PIQ purity score among all transcription factor motif instances and the target gene. Furthermore, the confidence score was converted to percentile ranks ranging from 0 to 1. Only edges with confidence score > 0.99 were preserved in the final network, containing 1,013 outgoing edges of *POU3F2*.

### *De novo* mutation enrichment analysis

We used published *de novo* mutability^61^ of synonymous, missense, and loss-of-function variants to estimate the expected counts of mutations. Published *de novo* mutation data^5^ in 2,508 probands and 1,911 controls from the SSC cohort were accessed through denovo-db^62^. Loss-of-function mutations were defined as frameshift, stop-gained, splice-donor, stop-gained near splice, frameshift near splice, stop-lost, or splice-acceptor mutations. Missense mutations included missense and missense-near-splice labels from the denovo-db. Synonymous mutations included synonymous and synonymous-near-splice labels. We used Poisson test to assess enrichment and quantify the statistical evidence^61^.

### Partitioned heritability analysis

We used stratified LDSC^36^ to assess the partitioned ASD heritability in *POU3F2* TFBS. We used the PIQ motif instances we generated in the network analysis and expanded each TFBS by 100, 150, and 250 base pairs up- and downstream. Further, we partitioned the heritability from the using the meta-analyzed GWAS summary statistics as input. The model also included 53 LDSC baseline annotations, as recommended in Finucane et al^36^.

### URLs

AGP (https://www.ncbi.nlm.nih.gov/projects/gap/cgi-bin/study.cgi?study_id=phs000267.v5.p2);

SSC (https://www.sfari.org/resource/simons-simplex-collection/);

SPARK (https://www.sfari.org/resource/spark/);

SFARI Genes database (https://gene.sfari.org/about-gene-scoring/);

SPARK Genes (https://simonsfoundation.s3.amazonaws.com/share/SFARI/SPARK_Gene_List.pdf);

Fetal brain Hi-C data (https://www.ncbi.nlm.nih.gov/geo/query/acc.cgi?acc=GSE77565);

BRAINSPAN (http://www.brainspan.org/static/home).

### Data and code availability

Summary statistics from the ASD GWAS and TWAS are freely accessible at (ftp://ftp.biostat.wisc.edu/pub/lu_group/Projects/TITANS). The code to perform trio-based TWAS and GWAS analysis is available at (https://github.com/qlu-lab/TITANS).

## Acknowledgements

We are grateful to all the families participating in the Autism Genome Project (AGP), the Simons Simplex Collection (SSC), and the Simons Foundation Powering Autism Research for Knowledge (SPARK) study. This project was supported by the Clinical and Translational Science Award (CTSA) program, through the NIH National Center for Advancing Translational Sciences (NCATS), grant UL1TR000427. We also acknowledge research support from the University of Wisconsin-Madison Office of the Chancellor and the Vice Chancellor for Research and Graduate Education with funding from the Wisconsin Alumni Research Foundation and the Waisman Center pilot grant program at the University of Wisconsin-Madison. We thank Drs. Jakob Grove and Elise Robinson for sharing the GWAS summary statistics based on the iPSYCH cohort. We thank Drs. Brittany Travers, James Li, Xinyu Zhao, Jan Greenberg, and Marsha Mailick for helpful discussions.

## Author contribution

Q.L. conceived and designed the study.

K.H. and Q.L. developed the statistical framework.

K.H., Y.W., Z.N., J.Y., and J.C. performed the statistical analysis.

Y.W. and K.H. conducted data processing and quality control.

S.R., J.S., and A.F.S. assisted in network analysis.

Y.Z. and S.K. assisted in Hi-C data processing.

D.W. assisted in the analysis of spatiotemporal gene expression data

K.H., Y.W., and Y.L. implemented the software.

Q.L. advised on statistical and genetic issues.

K.H. and Q.L. wrote the manuscript.

All authors contributed in manuscript editing and approved the manuscript.

