## Supplementary Figures for "Transcriptome-wide transmission disequilibrium analysis identifies novel risk genes for autism spectrum disorder"

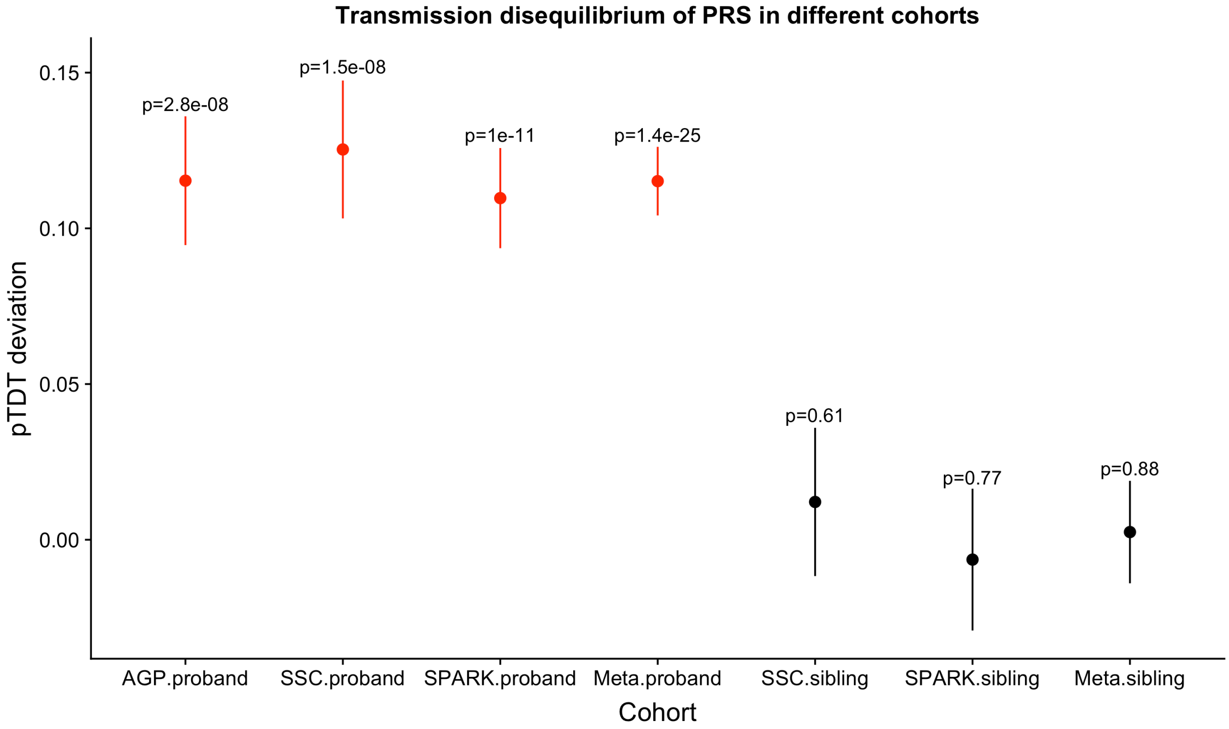


**Supplementary Figure 1. Transmission disequilibrium of PRS in different cohorts.** Transmission disequilibrium was quantified by the pTDT approach. Results in probands and unaffected siblings are highlighted in different colors. The mean pTDT deviation and the SE are shown. P-values are labeled above each interval.

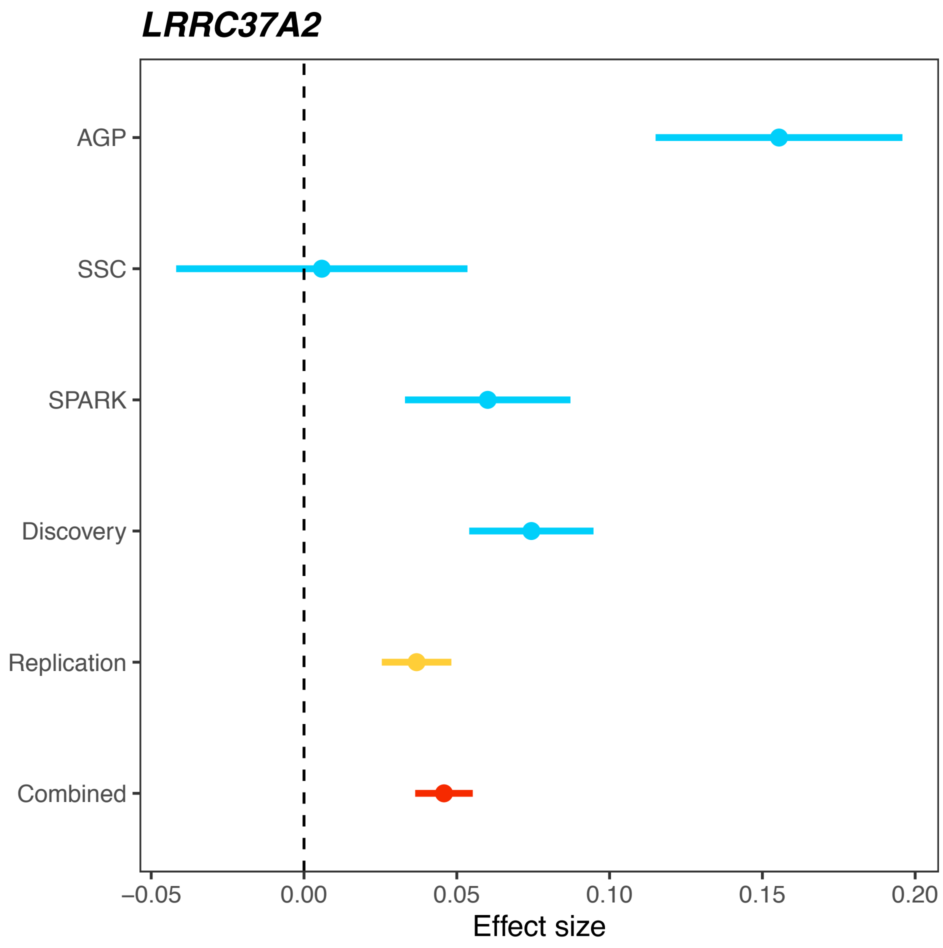


**Supplementary Figure 2. Forest plot for the significant association in GTEx anterior cingulate cortex BA24.** *LRRC37A2* reached transcriptome-wide significance in the TWAS in GTEx anterior cingulate cortex BA24. Standardized effect sizes (beta) and SEs are provided for all cohorts. Beta and SE in the discovery cohort are meta-analyzed results based on AGP, SSC, and SPARK. Beta and SE in the combined cohort are calculated from the meta-analysis of discovery and replication stages.

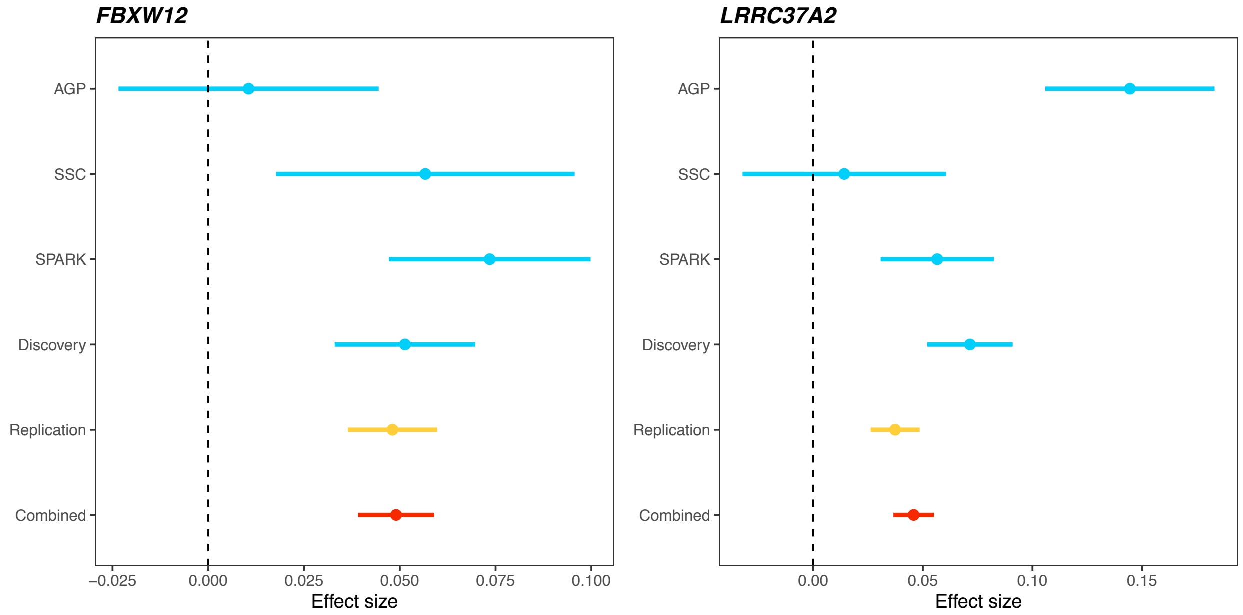


**Supplementary Figure 3. Forest plot for significant associations in GTEx caudate basal ganglia.** *FBXM12* and *LRRC37A2* reached transcriptome-wide significance in the TWAS in GTEx caudate basal ganglia. Standardized effect sizes (beta) and SEs are provided for all cohorts. Beta and SE in the discovery cohort are meta-analyzed results based on AGP, SSC, and SPARK. Beta and SE in the combined cohort are calculated from the meta-analysis of discovery and replication stages.

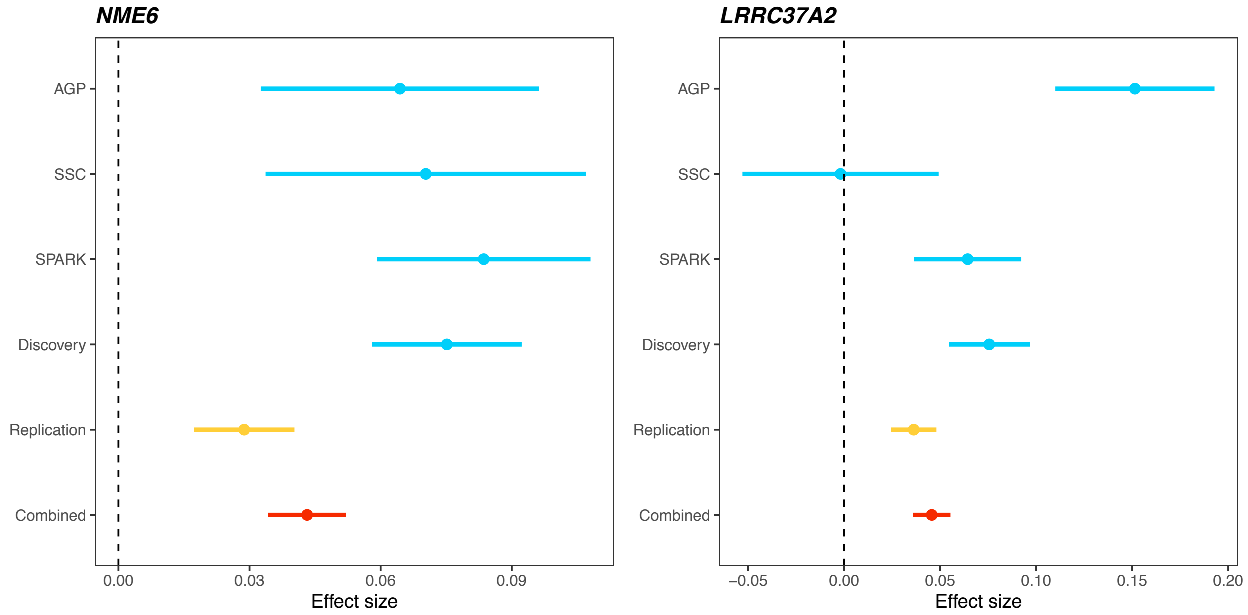


**Supplementary Figure 4. Forest plot for significant associations in GTEx cerebellar hemisphere.** *NME6* and *LRRC37A2* reached transcriptome-wide significance in the TWAS in GTEx cerebellar hemisphere. Standardized effect sizes (beta) and SEs are provided for all cohorts. Beta and SE in the discovery cohort are meta-analyzed results based on AGP, SSC, and SPARK. Beta and SE in the combined cohort are calculated from the meta-analysis of discovery and replication stages.

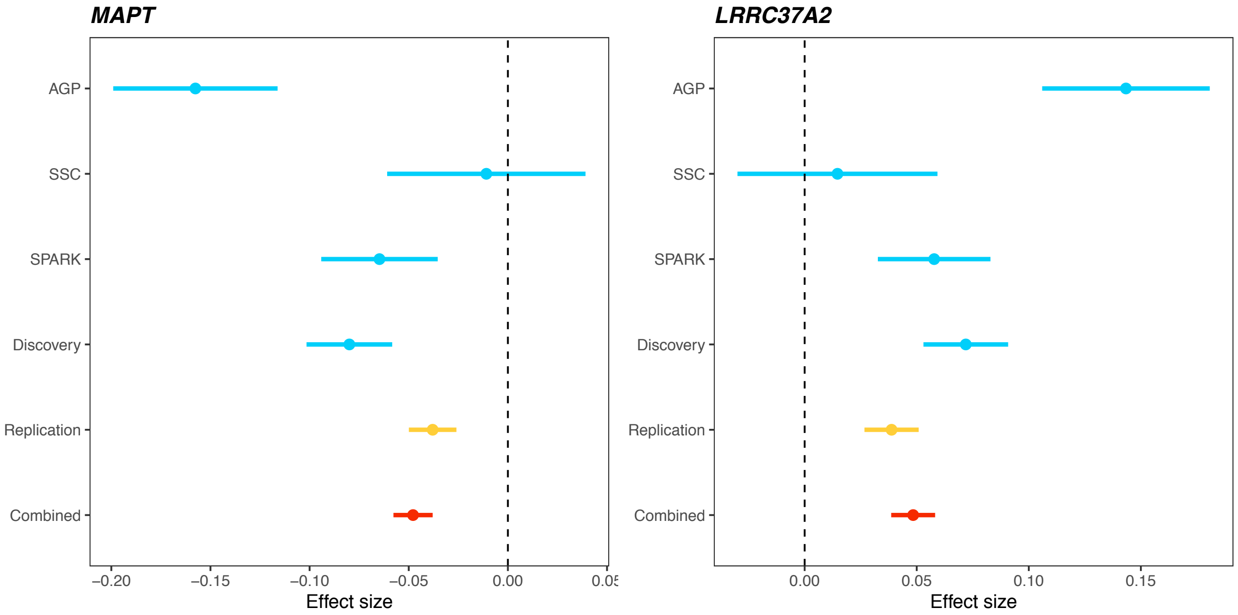


**Supplementary Figure 5. Forest plot for significant associations in GTEx cerebellum.** *MAPT* and *LRRC37A2* reached transcriptome-wide significance in the TWAS in GTEx cerebellum. Standardized effect sizes (beta) and SEs are provided for all cohorts. Beta and SE in the discovery cohort are meta-analyzed results based on AGP, SSC, and SPARK. Beta and SE in the combined cohort are calculated from the meta-analysis of discovery and replication stages.

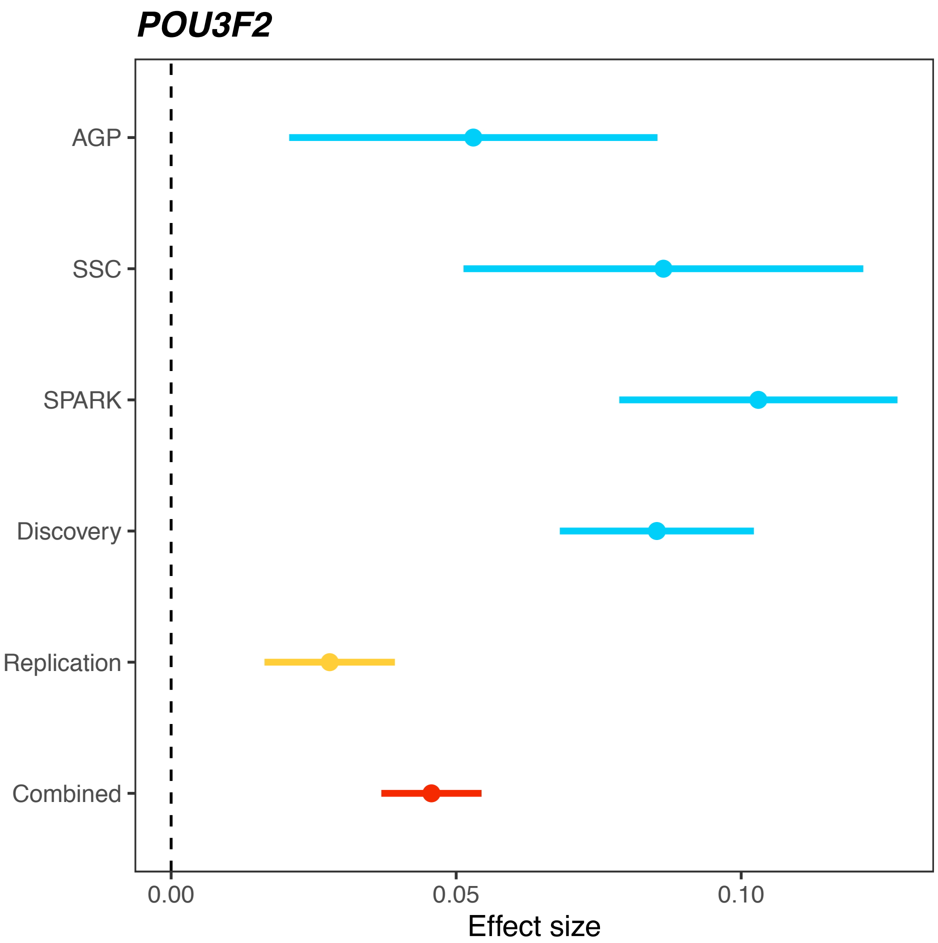


**Supplementary Figure 6. Forest plot for the significant association in GTEx hippocampus.** *POU3F2* reached transcriptome-wide significance in the TWAS in GTEx hippocampus. Standardized effect sizes (beta) and SEs are provided for all cohorts. Beta and SE in the discovery cohort are meta-analyzed results based on AGP, SSC, and SPARK. Beta and SE in the combined cohort are calculated from the meta-analysis of discovery and replication stages.

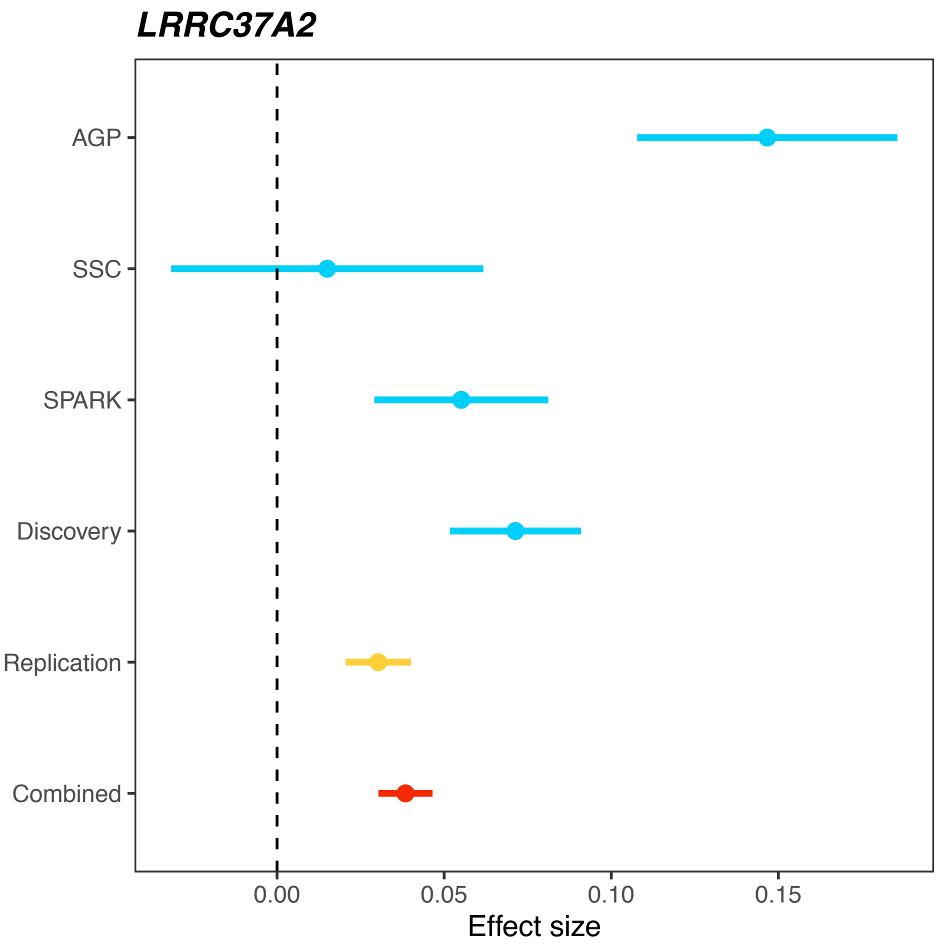


**Supplementary Figure 7. Forest plot for the significant association in GTEx hypothalamus.** *LRRC37A2* reached transcriptome-wide significance in the TWAS in GTEx hypothalamus. Standardized effect sizes (beta) and SEs are provided for all cohorts. Beta and SE in the discovery cohort are meta-analyzed results based on AGP, SSC, and SPARK. Beta and SE in the combined cohort are calculated from the meta-analysis of discovery and replication stages.

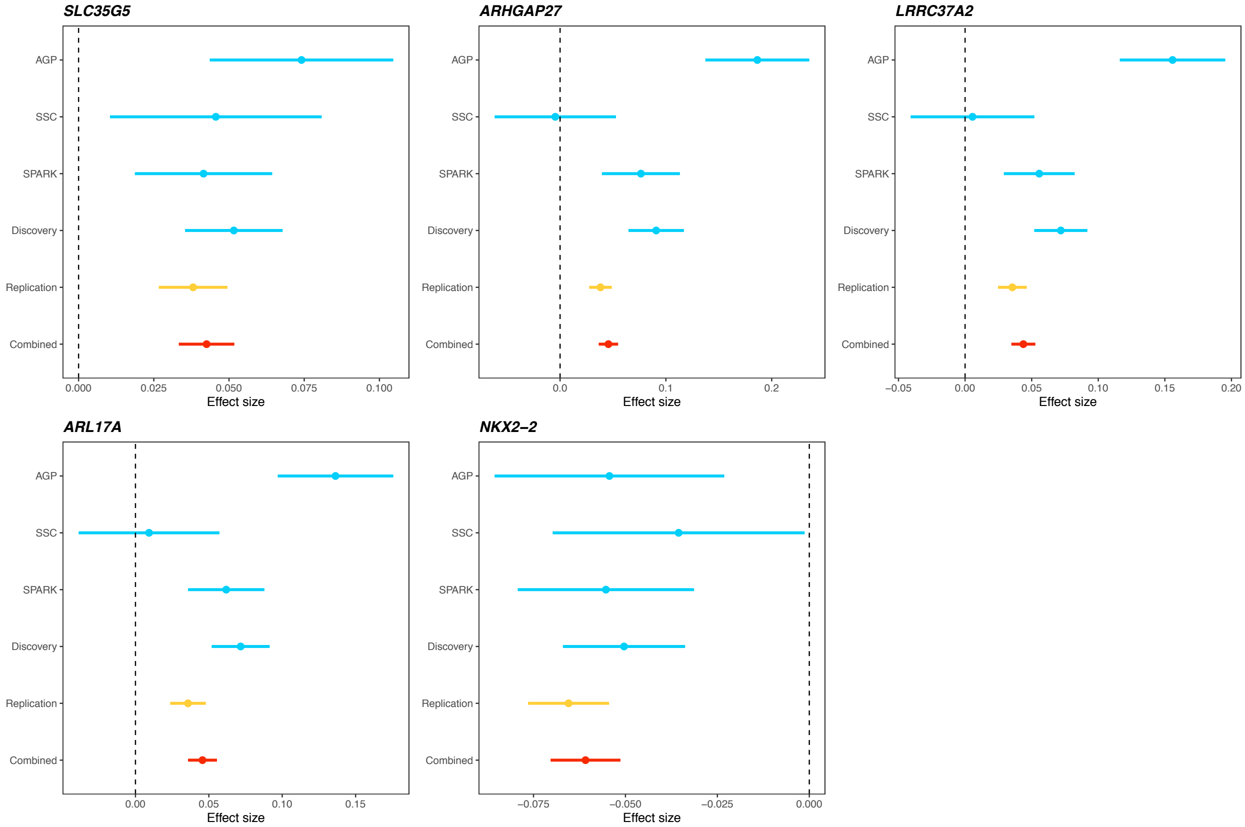


**Supplementary Figure 8. Forest plot for significant associations in GTEx nucleus accumbens basal ganglia.** *SLC35G5*, *ARHGAP27*, *LRRC37A2*, *ARL17A*, and *NKX2-2* reached transcriptome-wide significance in the TWAS in GTEx nucleus accumbens basal ganglia. Standardized effect sizes (beta) and SEs are provided for all cohorts. Beta and SE in the discovery cohort are meta-analyzed results based on AGP, SSC, and SPARK. Beta and SE in the combined cohort are calculated from the meta-analysis of discovery and replication stages.

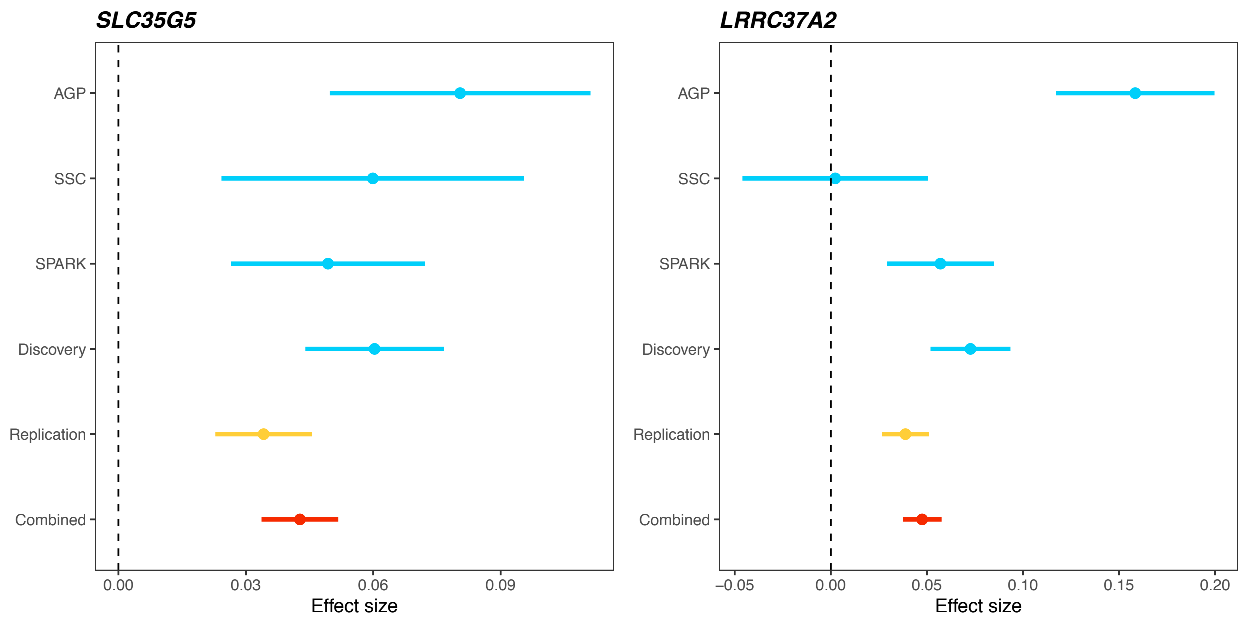


**Supplementary Figure 9. Forest plot for significant associations in GTEx putamen basal ganglia.** *SLC35G5* and *LRRC37A2* reached transcriptome-wide significance in the TWAS in GTEx putamen basal ganglia. Standardized effect sizes (beta) and SEs are provided for all cohorts. Beta and SE in the discovery cohort are meta-analyzed results based on AGP, SSC, and SPARK. Beta and SE in the combined cohort are calculated from the meta-analysis of discovery and replication stages.

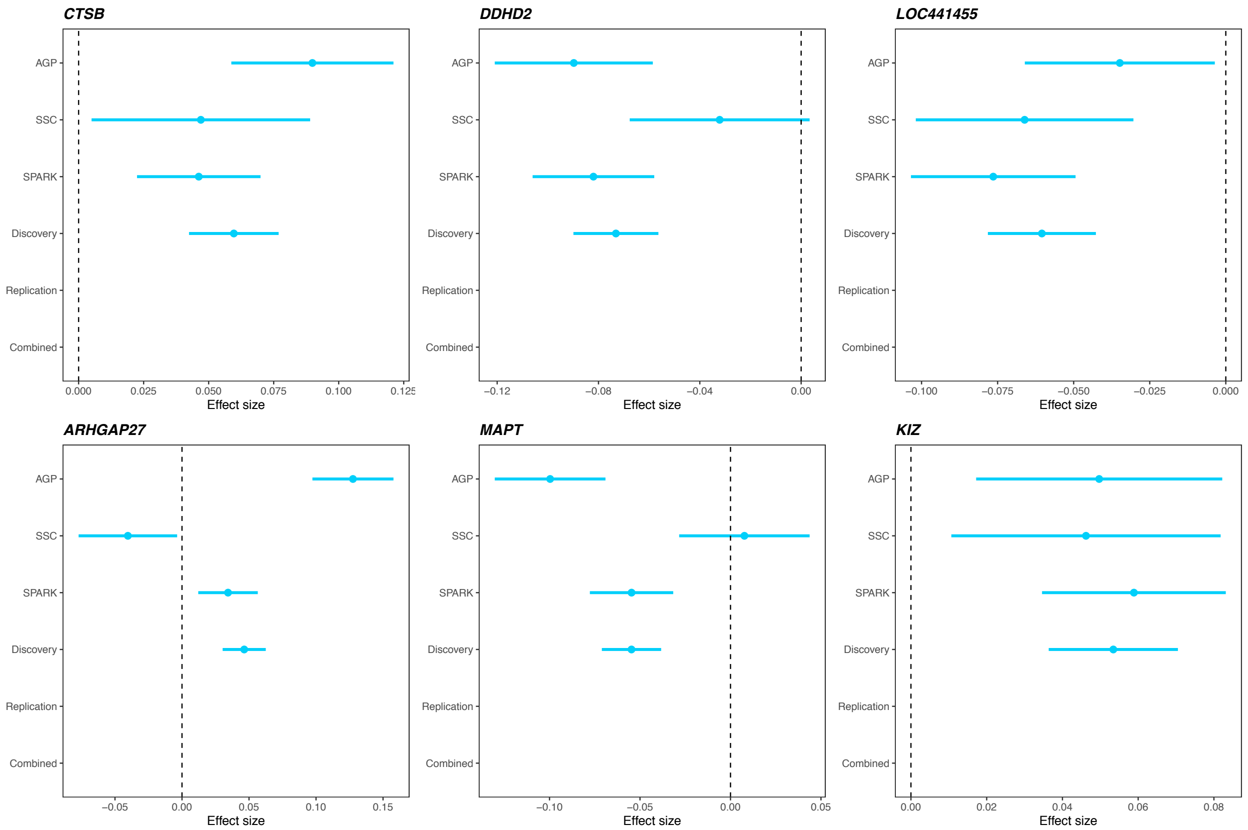


**Supplementary Figure 10. Forest plot for significant associations in CMC DLPFC.** *CTSB*, *DDHD2*, *LOC441455*, *ARHGAP27*, *MAPT*, and *KIZ* reached transcriptome-wide significance in the TWAS in CMC DLPFC. Standardized effect sizes (beta) and SEs are provided for the trio-based cohorts. Beta and SE labeled as the discovery cohort are meta-analyzed results based on AGP, SSC, and SPARK. Effect estimates are not shown in the replication and the combined cohorts since FUSION does not output beta and SE estimates.

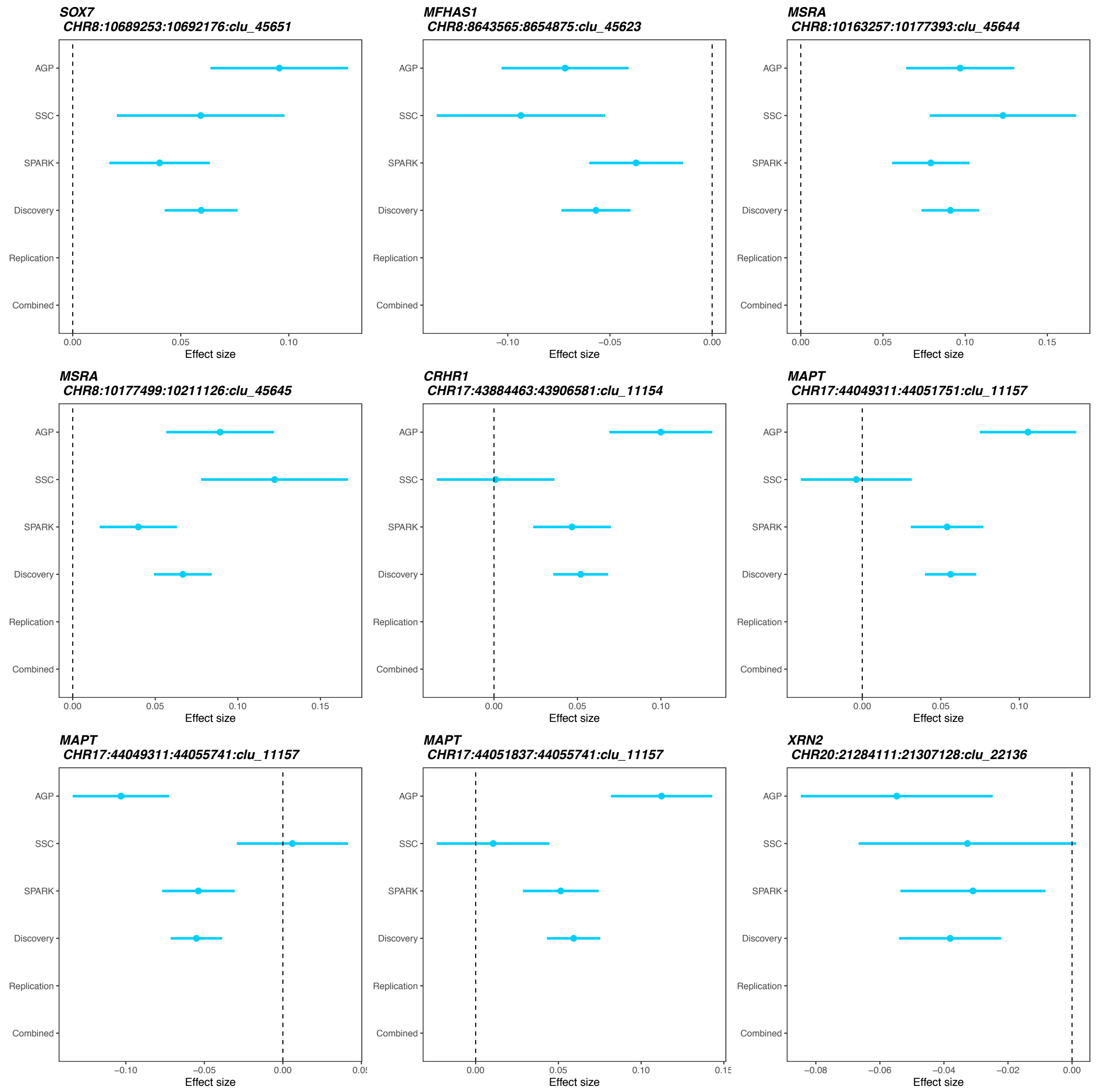


**Supplementary Figure 11. Forest plot for significant associations in CMC DLPFC splicing.** *SOX7*, *MFHAS1*, *MSRA*, *CRHR1*, *MAPT*, and *XRN2* reached transcriptome-wide significance in the TWAS in CMC DLPFC splicing. Intron cluster IDs are shown below the gene names. Standardized effect sizes (beta) and SEs are provided for the trio-based cohorts. Beta and SE labeled as the discovery cohort are meta-analyzed results based on AGP, SSC, and SPARK. Effect estimates are not shown in the replication and the combined cohorts since FUSION does not output beta and SE estimates.

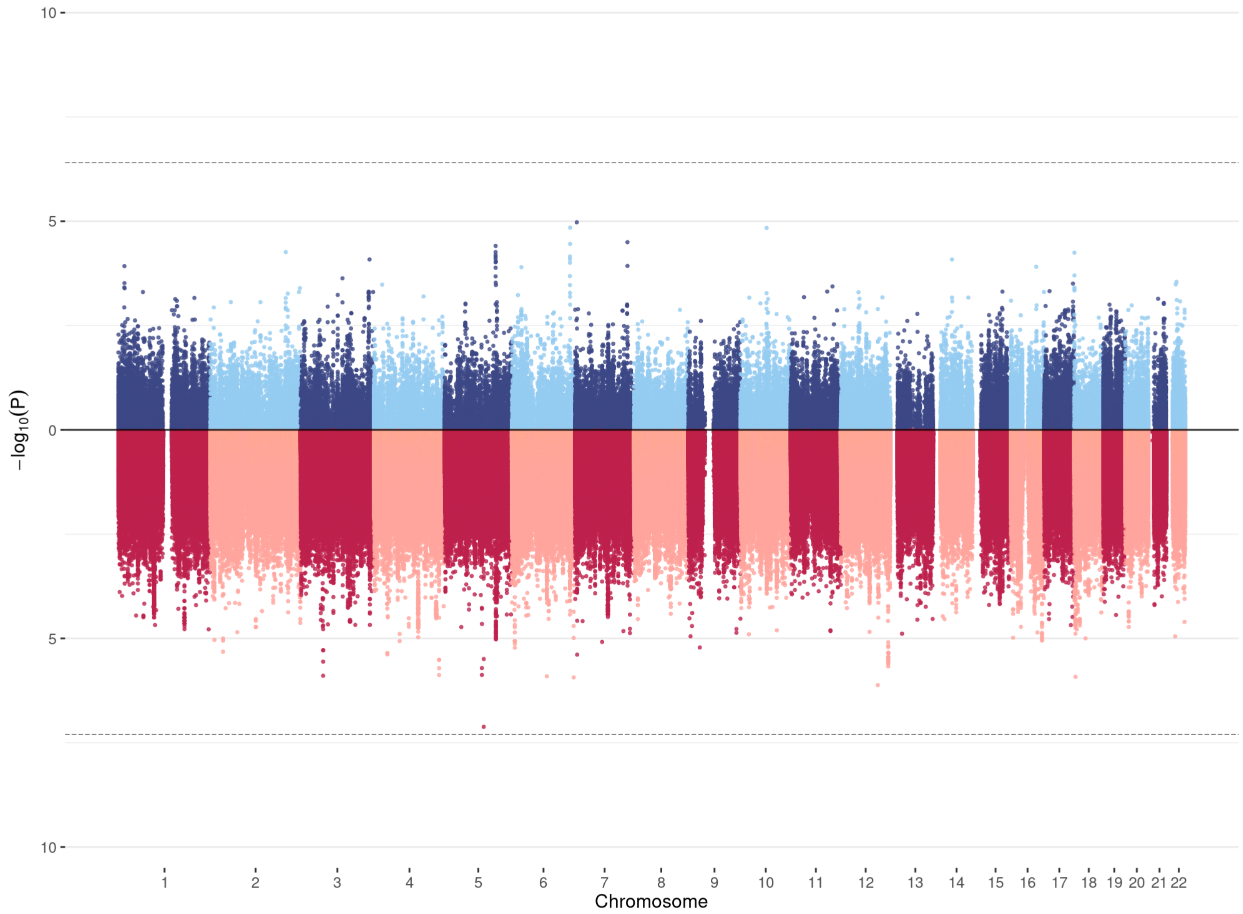


**Supplementary Figure 12. Mirrored Manhattan plot for TWAS and GWAS results in 3,245 sibling-parent trios.** TWAS results are shown in the upper panel. GWAS associations are shown in the lower panel. The dashed line in the upper panel indicates the cross-tissue transcriptome-wide significance cutoff (p=4.0e-7) and the dashed line in the lower panel is the genome-wide significance cutoff (p=5.0e-8). TWAS associations for all 12 tissues are shown.

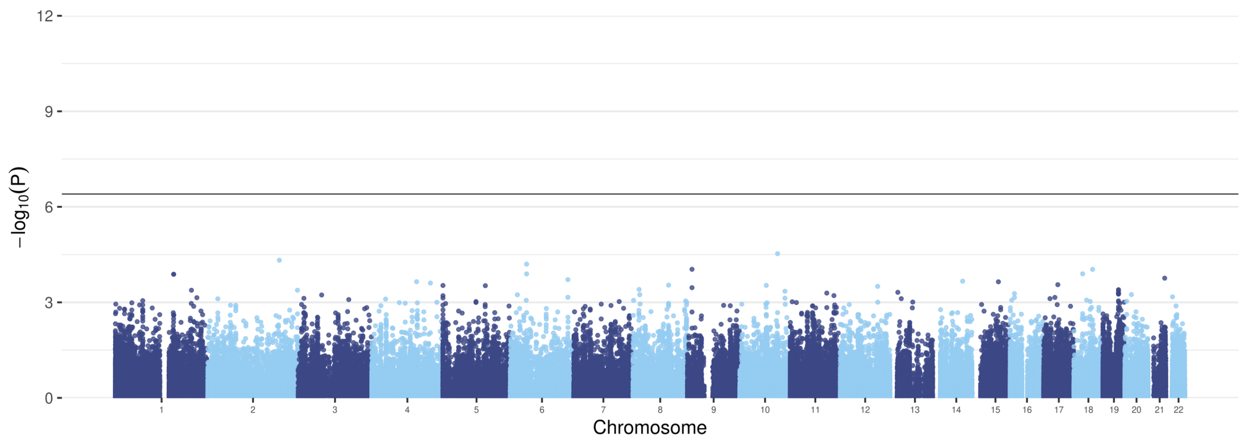


**Supplementary Figure 13. Manhattan plot for TWAS in 7,805 proband-parent trios after randomly shuffling the status of probands and pseudo siblings.** The dashed line indicates the cross-tissue transcriptome-wide significance cutoff (p=4.0e-7). TWAS associations for all 12 tissues are shown.

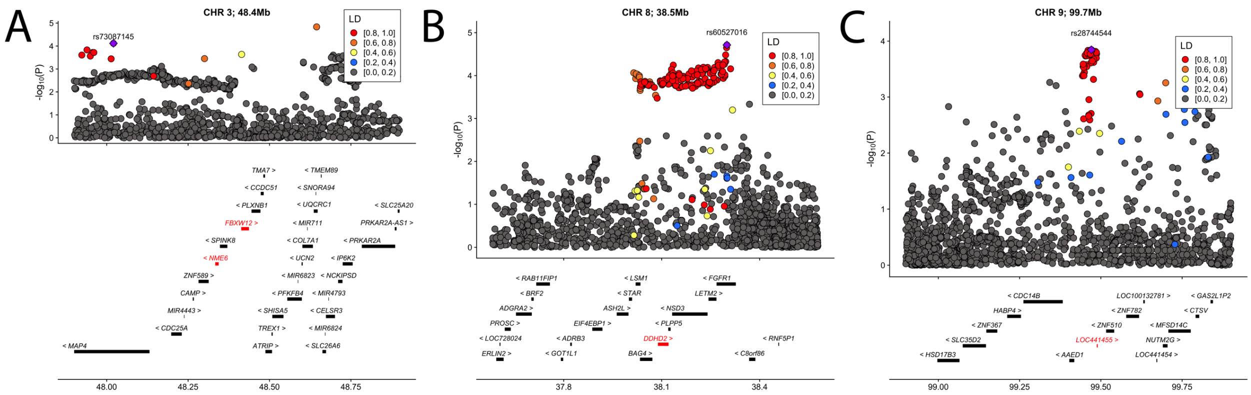


**Supplementary Figure 14. Additional significant loci identified in TWAS.** We identified 31 transcriptome-wide significant associations from 7 independent loci. Four loci with associations that remained significant after correcting for all genes and all tissues are shown in Figure 3 in the main text. **(A)** chr3, 48.4 mb **(B)** chr8, 38.5 mb **(C)** chr9, 99.7 mb. For each locus, the index SNP with the most significant association in GWAS is marked as purple diamond and the color of data points indicates LD of neighboring SNPs with the index SNP. Genes are highlighted in red if they reached transcriptome-wide significance in at least one tissue. The x-axis denotes genome coordinates and the y-axis denotes association p-values in GWAS.

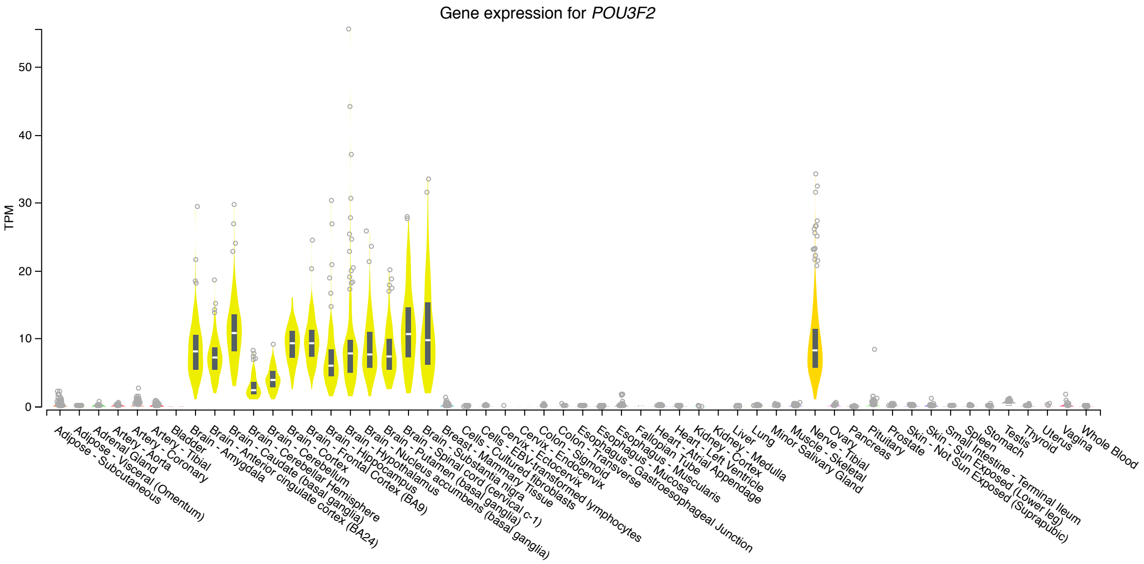


**Supplementary Figure 15. Multi-tissue gene expression profile of *POU3F2* in GTEx Release V8.**

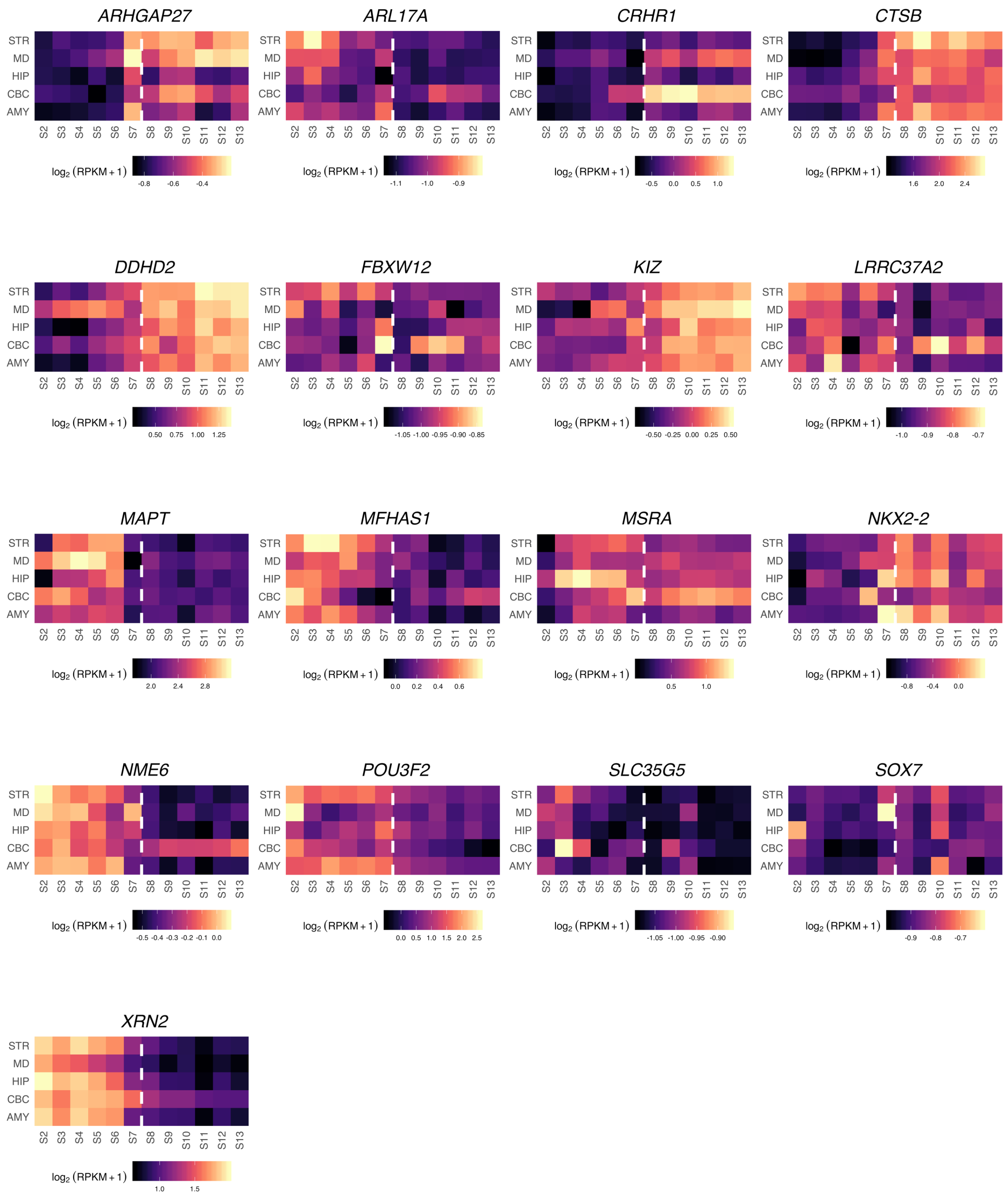


**Supplementary Figure 16. The spatiotemporal expression pattern of candidate genes identified in TWAS**. The spatiotemporal expression pattern of 17 TWAS genes across 5 brain regions and 12 developmental stages. The dashed line indicates the boundary between later fetal and early infancy stages (0 month).
